# Mega-frequency mutagenesis: generation of non-random precise mutations with extremely high frequency upon adaptation of cancer cells to drugs and stress

**DOI:** 10.64898/2026.02.20.707073

**Authors:** Victor Oleynik, Anjana Edathil Kadangodan, Valid Gahramanov, Subha Ranjan Das, Bar Levi, Julia Yaglom, Kirill Anoshkin, Santosh Kumar, Benjamin G. Steinberg, Yitzhak Reizel, Pinchas Polonsky, Igor Koman, Vadim Levitt, Albert Pinhasov, Andriy Marusyk, Elimelech Nesher, Michael Y. Sherman

**Author notes:** To whom correspondence should be sent. Equal contribution (Victor Oleynik, Anjana Edathil Kadangodan).

## Abstract

Acquiring drug resistance is a major problem in cancer treatment. As cancers adapt to chemotherapy, chromatin landscape becomes altered in subpopulations of persister cells to acquire gene expression patterns that provide drug resistance. The increased level of stress-induced random mutagenesis in cancer has also been linked to acquisition of drug resistance. Here we show that during adaptation to conventional cytotoxic chemotherapies or targeted therapies, tens of thousands of mutations are generated, and the probability of acquisition of these mutations at specific positions can reach 50% and even higher. A large fraction of these mutations is highly recurrent and non-random. The patterns of the recurrent mutations are specific to the drug target and are unrelated to the chemical nature of the drug. Surprisingly, these mutations are progressively generated at the non-dividing pseudo-senescence stage following drug exposure, and at this stage, selection is not involved in their accumulation. Notably, these mutations are highly enriched within or near binding motifs of certain transcription factors, like KLF9, IRF1 and others. Therefore, a mechanism for precise generation of mutations at specific positions with extremely high rates appears to be triggered upon drug adaptation, and these precise mutations may affect activities of a set of transcription factors.

## Introduction

Development of drug resistance limits the effectiveness of cancer therapies. This phenomenon is especially evident with targeted therapies, e.g. tyrosine kinase inhibitors that induce strong clinical responses, ultimately followed by relapse. For example, pioneering work with inhibition of B-Raf showed almost complete disappearance of melanoma metastasis, followed by their re-appearance several months later (1-3). Therefore, targeted therapies typically fail to eradicate all tumor cells, and residual cells acquire resistance causing cancer relapses. There is extensive literature on mechanisms of drug resistance, such as overexpression of MDR pumps, mutations in drug targets, activation of drug-metabolizing enzymes, and others (4-8). However, in contrast to detailed elucidation of specific molecular mechanisms underlying drug resistance, investigation of the origin of resistant mutations received much less attention.

Recent studies with cell cultures uncovered that resistance emerges in the process of drug treatment from preexistent drug-tolerant persister cells that show increased levels of expression of protective genes [9] and gradually acquire additional transcriptome changes to achieve fully resistant phenotypes (e.g. to ALK inhibitors) (10,11). Persisters initially represent weakly resistant heterogeneous subpopulations, which differ in fitness when exposed to different tyrosine kinase inhibitors, for example (12,13). Resistance gradually increases during therapy through acquisition of multiple epigenetic drug-specific mechanisms. For example, resistance to EGFR inhibitors such as erlotinib develops gradually and depends on the histone methylation status (14-16). In line with these observations, development of resistance to a series of EGFR and ALK inhibitors was associated with changes in the patterns of chromatin opening (11). It was found that in the process of the resistance development, different subpopulations of cells transit through a set of relatively fixed gene expression patterns, evolving from one pattern to another (11).

Besides epigenetic mechanisms, genetic changes also appear to play a role in resistance development. Indeed, tracking the fates of drug-treated cells using DNA barcoding demonstrated that in some cases, resistance arises due to selection of pre-existent resistant mutant subpopulations (17), which was in line with a paradigm that development of resistance to anti-cancer therapies is due to mutations that exist prior to treatment (18,19). Accordingly, relapses are thought to occur because drug-resistant mutant cells are present in primary tumors and metastases prior to initiation of therapy, can survive drug treatments and enable recovery from the treatment. However, recent studies demonstrate that resistance to targeted therapies can also be facilitated by a transient increase in genomic instability at the time of treatment, leading to generation of de novo mutations. A similar process can lead to antibiotic resistance in microbes, where stressful treatments induce genetic instability and increase mutation rates in bacteria and yeast (20-23). The increased mutability was associated with the reduction in the efficiency of DNA mismatch repair (24,25), and upregulation of error-prone DNA polymerases (26-29). It was argued that once the cells became adapted to stress, the mutation rates subsided due to a counter-selection against accumulation of deleterious mutations (21,30,31).

In cancer studies, it was demonstrated that upon exposure to various therapies, cells downregulate genes of the mismatch repair and homologous recombination repair systems, which lead to an increased mutation rate (32-34). It was suggested that the elevated rate of mutations can facilitate the development of resistance.

One of the mechanisms of generation of mutations involves APOBEC enzymes that catalyze deamination of cytosines in DNA, thus creating transition from C to T or G. High expression of APOBEC was associated with increased drug resistance in colon and breast cancer, while inactivation of APOBEC3A significantly impairs the development of drug resistance (35-37). These data were interpreted within a conventional genetic framework as indication of the role of the elevated random mutagenesis in acquisition of the drug-resistance mutations that are fixed in the cell population by selection for resistant variants.

Here, we sought to further explore drug-induced generation of mutations. Unexpectedly, we uncovered that the mutational process upon exposure to drugs or stresses is not random. Indeed, these treatments trigger generation of thousands of identical mutations in different cells in the population in a treatment-specific manner, and remarkably, selection is not involved in accumulation of these mutations. These data indicate the existence of a mechanism for an active generation of mutations at specific locations in response to stressful treatments.

## Results

### Experimental design

To systematically study mutational processes during adaptation to environmental and therapeutic stressors, we examined de novo mutations that arise during adaptation to a spectrum of stressors characterized by distinct toxicities and proximal resistance mechanisms. Specifically, we chose doxorubicin (Top2 inhibitor), irinotecan (Top1 inhibitor), glucose deprivation (stress, not a drug); osimertinib (EGFR inhibitor), crizotinib and lorlatinib (ALK inhibitors) (see Table 1 for corresponding cell lines). To attain genetic homogeneity of cell populations, we isolated single clones of each cell line, which we used as PARENTAL CLONES in each experiment. This genetic homogeneity of each parental clone ensures that mutations that arise during drug treatment are generated in the course of the experiment and did not preexist. Treatments started when the number of cells in parental clones reached 2mln, which corresponded to approximately 21 cell divisions. Notably, the rate of spontaneous generation of mutations in the untreated parental cells was incomparably lower than in drug-treated cells (see experiment with Alk inhibitors, below, where 13,737 mutations were generated in a clone of lorlatinib-treated cells and 842 in untreated parental cells).

**Table 1.** Cell lines used in experiments with different drugs and stress.

| Treatment | Target | Cell line | Cancer type |
| --- | --- | --- | --- |
| Doxorubicin | Top2 | HCT116 | Colon |
| Irinotecan | Top1 | HCT116 | Colon |
| Osimertinib | EGFR | H1975 | Lung |
| Lorlatinib | ALK | H3122 | Lung |
| Crizotinib | ALK | H3122 | Lung |
| Glucose deprivation |  | HCT116 | Colon |

In the first type of experimental setup (setup 1), a single clonal population of cells was initially subjected to a drug for 24 hours at concentrations that result in the death of about 70-80% of the population within 72 hours (Fig. 1a). With all tested drugs, within two to three days, the surviving cells underwent growth arrest and entered a pseudo-senescent state akin to senescence. Cells had an enlarged and flattened morphology, were highly vacuolized and expressed senescence-associated β-gal (Fig. 1b, c). They remained in this state for weeks or even months, depending on the original drug concentration, and then a fraction of cells began dividing again, suggesting that they were able to adapt to the drug while being in the pseudo-senescent state (Fig. 1d). At subsequent treatments with the same drug concentration, the time periods that the cells remained in pseudo-senescent state got shortened gradually (e.g. with 12μM osimertinib, Fig. 1d). At further treatments, the doses of the drug were gradually increased to achieve resistance to higher doses (Fig. 1d), and following several cycles of treatment, cells developed resistance to the highest drug concentration.

**Fig. 1.**
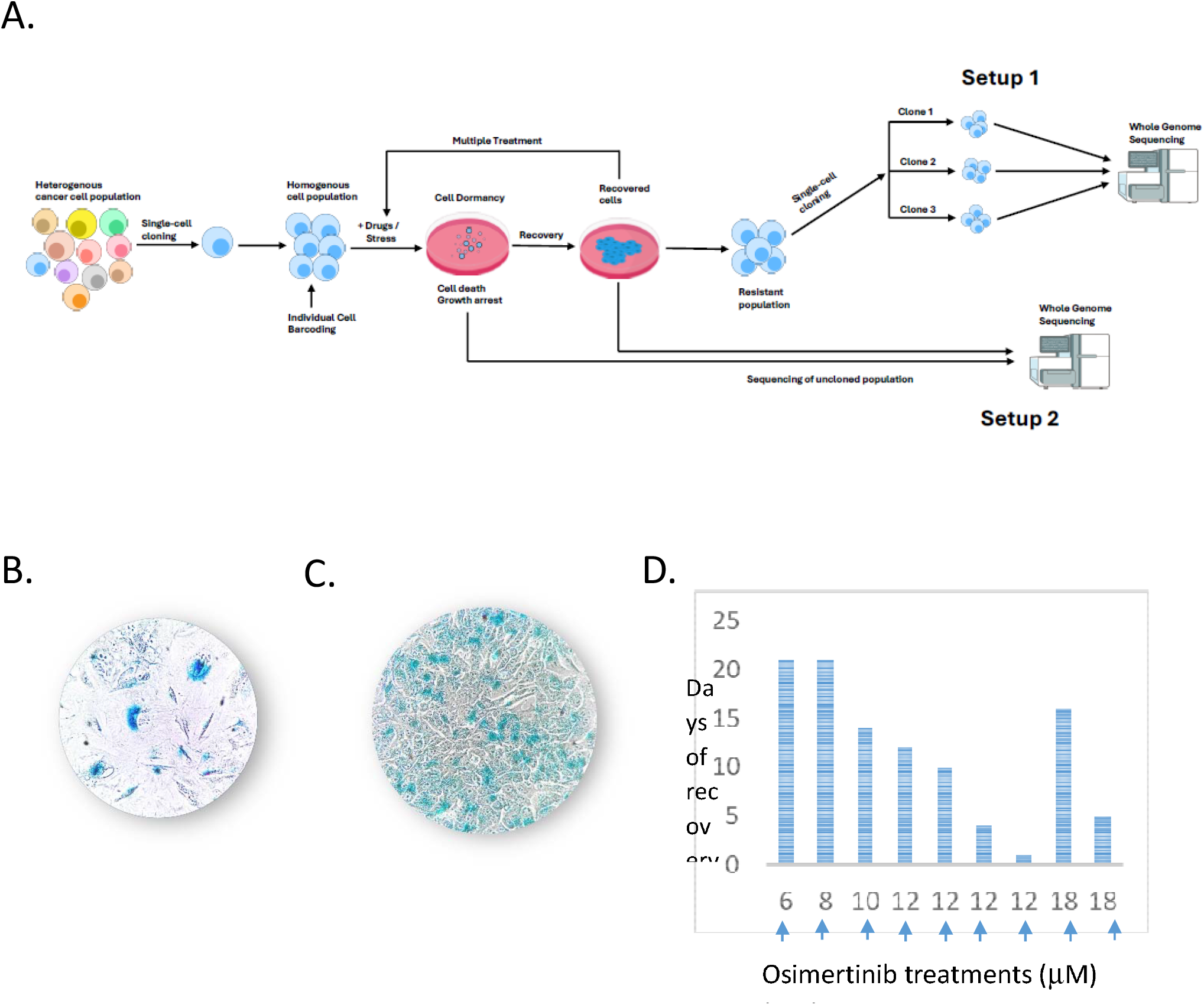
The experimental setup. A. Scheme of the experiment. B. Development of a temporal senescence (pseudo-senescence) in a population of osimertinib-treated cells. H1975 cells were treated with 6mM of osimertinib for 24h. On day 7 post treatment the cells were stained for the senescence-associated β-gal activity. C. Development of a temporal senescence in a population of Dox-treated cells. HCT116 cells were treated with 200nM of Dox for 24h. On day 5 post treatment the cells were stained for the senescence-associated b-gal activity. D. Kinetics of adaptation of cells to osimertinib (days in pseudo-senescence before the recovery). Cells at 50% confluence were treated with osimertinib for 24 h, and the drug was omitted. A fraction of cells died with the next three days and the remaining cells persisted in pseudo-senescent state. The number of days before cells started dividing and filled the plate was measured. On X-axis, arrows indicate doses of osimertinib in sequential treatments with increasing concentrations of the drug.

Importantly, prior to drug treatments, cells of the expanded parental clone were individually DNA barcoded. After achieving the resistance, cells were subcloned again, and barcodes in individual subclones were sequenced. The presence of distinct barcodes in the selected drug-resistant subclones ensured that they are descendants of cells that diverged before the treatments, and thus all mutations are generated in different cells independently. Cells that developed resistance to drugs were cloned again (Fig. 1a, setup 1), and two or three clones with different barcodes (for each drug treatment) were chosen for sequencing.

In a different experimental setup (setup 2), we administered the drug just once (e.g. for 24h with osimertinib or exposed continuously with Alk inhibitors) and did not do cloning of the recovered cells (only initial cloning to generate the parental clone). In these experiments, DNA for sequencing was isolated from the entire populations of pseudo-senescent or recovered cells (Fig. 1a, setup 2), which allowed tracking the dynamics of generation of mutations upon treatments.

### Emergence of location-specific mutations

Following whole genome sequencing in the setup 1 experiments, the genomes of three independent subclones resistant to a drug were compared to the genome of the parental clone, unveiling large numbers of mutations generated in the course of the treatments. Since our goal was to identify mutations with very high confidence, in calling mutations with Mutect2 package, along with the default mutation filtering, we used additional strong filtering parameters, including (a) no less than 20 good reads for a position that acquires a mutation, (b) in parental clone zero reads among both good and bad reads that carry mutant nucleotide in a given position, (c) the presence of at least 20% of reads with the mutant nucleotide in a given position in resistant clones (Table 2, Table S1). In setup 2, since the sequencing was done with a mixed (not cloned) population of cells, we expected a lower fraction of sequencing reads that carry mutations (some cells carry a particular mutation and some do not). Therefore, the filters applied upon mutation calling were slightly different, (a) no less than 20 good reads for a position that acquires mutation, (b) in parental clone zero reads among both good and bad reads that carry mutant nucleotide in the position, (c) the presence of at least 10% of reads with the mutant nucleotide or at least 3 mutant reads with mutant nucleotide in the position in resistant cells, whatever is higher. In the analysis of mutations, we did not consider *de-novo* InDels or loss of heterozygosity (LOH). The main reason was that in irinotecan or doxorubicin-treated cells, de novo InDels or LOH (unlike SNPs) originate from double strand DNA breaks at the sites of action of Top1 or Top2 (38, also manuscript in review). Therefore, with all treatments, we focused on de novo SNPs.

**Table 2.** Main steps of filtering upon mutation calling.

| Stage | Step Description | Purpose |
| --- | --- | --- |
| 1. Control coverage requirement | Retain only positions with sufficient read depth ($\geq 20$ reads) in the control sample. | To ensure confident assessment of reference genotype in the control. |
| 2. Reference allele validation in control | Select positions with $\geq 20$ reference reads and 0 alternative reads in control. | To guarantee no evidence of the alternative allele in the control; helps define high-confidence *de-novo* candidate sites. |
| 3. Remove weak sites | Extract positions with at least 3 alternative reads in at least one treated sample. | To eliminate likely sequencing noise. |
| 4. Separate mutation types | Extract SNPs. | To focus downstream analysis on SNPs. |
| 5. Validation against control BAM | Eliminate positions with alternative reads in control (parental clone) BAM file. | To verify *de-novo* status of SNPs in resistant cells files beyond VCF-based evidence. |
| 6. Clone-specific variant selection | Extract SNPs with alternative allele frequency $\geq 20\%$ in experimental setup 1 and $\geq 10\%$ in experimental setup 2. | To identify mutations within each treated sample, relative to control. |
| 7. Common mutation filtering | From stage 5, select variants detected in all treated samples with allele frequency $\geq 20\%$ in experimental setup 1 and $\geq 10\%$ in experimental setup 2. | To identify common mutations between treated samples. |

Remarkably, when we compared SNPs generated upon doxorubicin treatment, more than 80% of them (11,806 mutations) were identical in all three independently treated subclones (Fig. 2a). In another example, we observed that 12% of the total SNPs (402 mutations) were identical in three independent osimertinib-resistant subclones (Table 3). Similarly, multiple identical mutations were seen in independent subclones with all tested treatments (Tables 3,4). Therefore, it appears that drug treatment of cells leads to independent generation of identical mutations.

**Table 3.** Comparison of mutations between different subclones of cells adapted to DOX, Iri and Osi.

| TREATMENT | SUBCLONE | TOTAL MUTATIONS | COMMON DOUBLE MUTATIONS |  | COMMON MUTATIONS | % of total | p-value |
| --- | --- | --- | --- | --- | --- | --- | --- |
| OSIMERTINIB<br>9 cycles<br>H1975 cells | 1 | 3469 | Clones 1-2 | 98 | 402 | 11.59 | $7 \times 10^{-468}$ |
|  | 2 | 6241 | Clones 1-3 | 218 |  |  |  |
|  | 3 | 5311 | Clones 2-3 | 132 |  |  |  |
| IRINOTECAN<br>23 cycles<br>HCT116 cells | 1 | 13322 | | | 1165 | 8.74 | $3 \times 10^{-255}$ |
|  | 2 | 13912 |  |  |  |  |  |

**Table 4.**
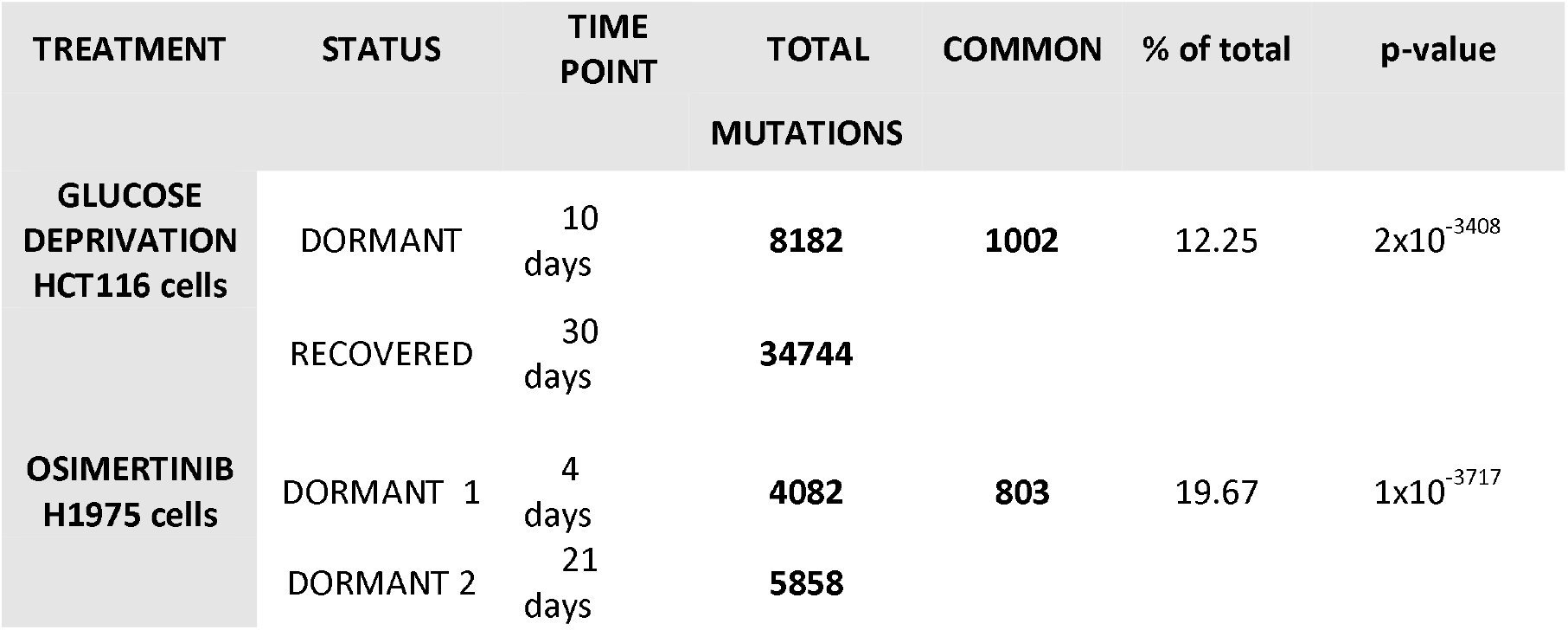
Comparison of mutations between different plates following treatment with Osi at days 4 and 21 of dormancy. Comparison of mutations between different plates following glucose deprivation at days 10 of dormancy and at day 30 upon the recovery.

**Fig. 2.**
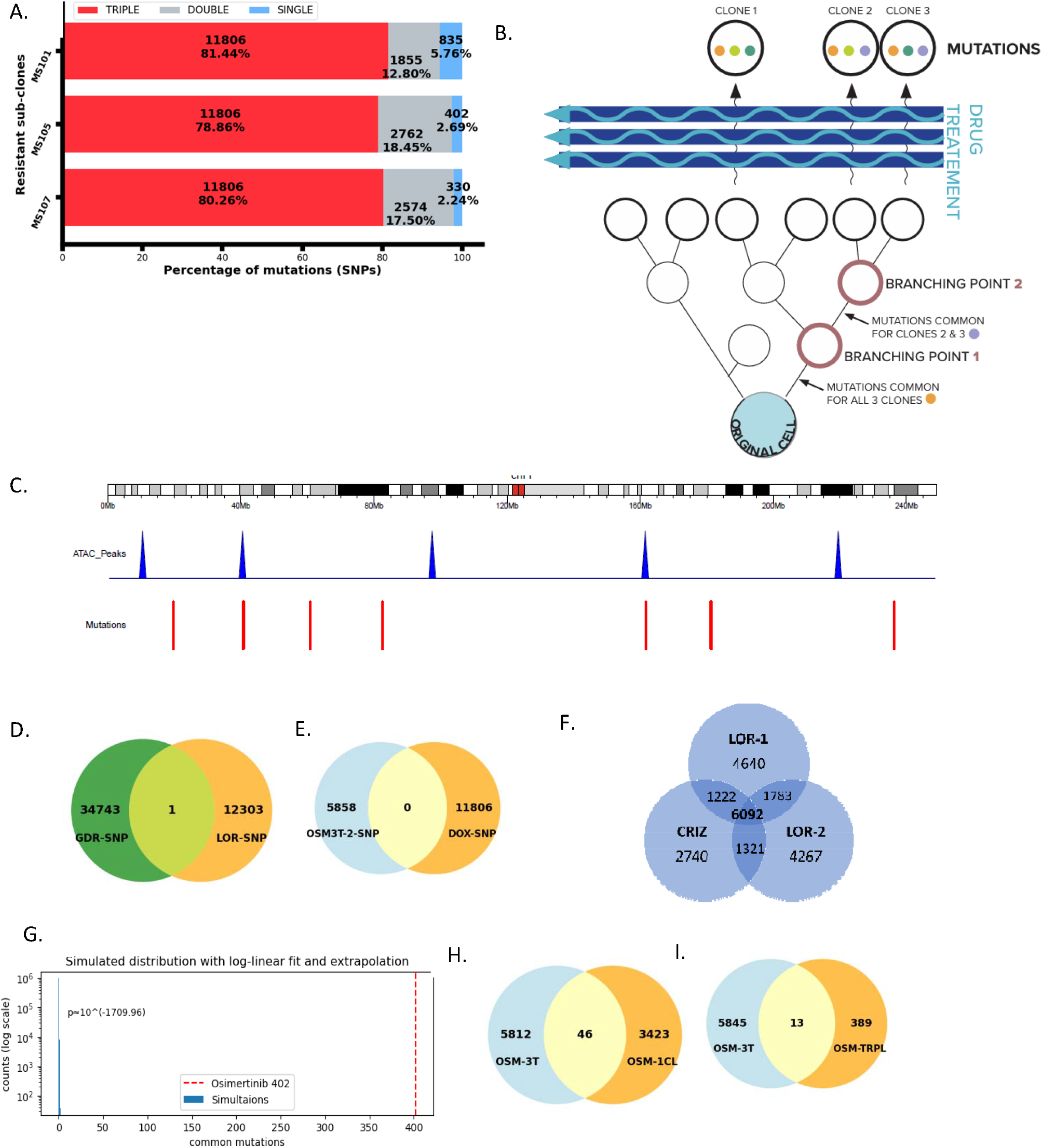
Development of recurrent mutations after drug treatments. A. The number of mutations in three parallel subclones following development of resistance to Dox. The number of mutations common to all three subclones (red), common to two out of three subclones (green) and mutations unique for each subclone (blue). B. A scheme showing that generation of mutations common between clones cannot occur during the propagation of the parental clone. See explanation in the text. C.Example of the lack of enrichment of mutations in open chromatin. Open chromatin assessed by ATACseq is shown in the lane with blue peaks, positions of mutations are shown in the lane with red bars. Data analysis in this experiment is shown in Table S2. C. Overlap of mutations between GDR and Lorlatinib. D. Overlap of mutations between osimertinib and Doxorubicin. E. Overlap of mutations between two independent lorlatinib-treated samples and one crizotinib-treated sample. F. Computer simulation of random generation of osimertinib-induced mutations. Observed number of common mutations is shown as red line. Random sampling of mutations and their overlap (100,000 pairs) is shown as blue lines. G. Overlap of mutations between osimertinib day 21 (0SM-3T) and osimertinib subclone 1 samples (OSM-1CL). H. Overlap of mutations between osimertinib day 21 (0SM-3T) and osimertinib mutations common between three subclones (OSM-TRPL).

An alternative model to explain appearance of the recurrent mutations in resistant subclones is that during the expansion of the parental clone from one cell to 2mln cells multiple mutations are generated spontaneously. A small subpopulation of cells from the expanded clone which carries a set of mutations survives a drug treatment. Since we chose our resistant subclones from this small subpopulation, they carry similar mutations. However, this model is improbable for several independent reasons. Firstly, since cells divide in a dichotomic branching manner, upon expansion of the parental clone, there must be two points of branching, one that separates one resistant subclone (clone 1) from two other resistant subclones, and another later branching point that separates two remaining resistant subclones (clones 2 and 3) (Fig. 2b). Accordingly, if the detected mutations are generated upon expansion of the parental clone, one can expect that three resistant subclones can have a number of mutations common for all three of them that were generated before the first branching point. Furthermore, subclones separated at the second branching point will have mutations common to two of them but not to subclone 1, which were generated during divisions between the first and the second branching points. However, there cannot be mutations that are common to subclone 2 and subclone 1, which are not shared with subclone 3. Similarly, there cannot be mutations that are common to subclone 3 and subclone 1, which are not shared with subclone 2 (Fig. 2b). We, however, see significant numbers of mutations that are shared by two out of three subclones, and these numbers are comparable in all three combinations of subclone pairs (Table 3, Fig. 2f). Therefore, common mutations must be generated in the process of drug treatment and cannot preexist and be selected.

Secondly, in a drug escalation experiment, cells undergo a much higher number of divisions than the number of divisions upon the expansion of the parental clone (around 21 divisions to reach 2mln cells starting from a single cell). In case of doxorubicin treatment there must be more 40 divisions in the dose escalation experiment, i.e. 20 cycles of treatment, and at least two divisions in each cycle to restore the number of cells in plates following initial cell death. Furthermore, drug treatments cause DNA damage and therefore are mutagenic. Accordingly, most of the mutations seen in resistant subclones must be generated upon drug treatments, and therefore it is impossible that following doxorubicin treatments 80% of mutations (which we see as common mutations) could result from selection of preexistent mutations in parental clone, rather than generated during drug treatments. Finally, as shown below, common mutations could be generated at the pseudo-senescent state in the absence of selection. Therefore, recurrent mutations must be generated in the process of drug treatments and not be selected from preexistent mutations in the parental clone. This very counterintuitive result raises the question why such a mutation recurrency has not been observed in WGS experiments previously, which is thoroughly addressed in the Discussion section.

Previous publications indicated that in some cancers, mutations are enriched in open chromatin (39), probably due to its vulnerability to stress, while in other cancers mutations are depleted in open chromatin, probably due to enhanced recruitment of the repair enzymes (40). To address whether the identified mutations associate with the open state of chromatin, we performed an ATACseq experiment with cells exposed to irinotecan or glucose deprivation. The state of the chromatin was monitored in cloned parental, pseudo-senescent and recovered cells. The results show that there was no enrichment of mutations in open chromatin, but rather a minor decrease (reduced probability of mutations falling in a peak) (Fig. 2c, Table S2). Accordingly, generation of mutations upon drug or stress treatments is not associated with the open state of chromatin.

### Drug target specificity of the mutation patterns

Analysis of mutations showed that their patterns were drug-specific (Fig. 2d,e), as almost no SNPs were common for different drugs. Notably, this drug-specificity further rejects the possibility the the mutations represent a technical artifact. An important insight regarding the drug-specificity of common mutations was obtained from cells treated with tyrosine kinase receptor inhibitors crizotinib and lorlotinib using H3122 cells. Structurally, these compounds are very distinct, while sharing the same major targets tyrosine kinase receptors Alk and ROS1. Analysis of SNPs generated in the presence these drugs showed that almost 60% of mutations (7,875) were identical between two independent parallel cultures treated with lorlatinib, and a similar fraction of identical SNPs was found between crizotinib and any of the lorlotinib samples (Fig. 2f). Notably, in this experiment in a control (DMSO-only) cell population that was passaged for 20 days (the time necessary to achieve recovery from ALK inhibitors), 842 mutations were seen, representing a low background level of drug-independent mutations. Even this comparably low number is a significant overestimation, since while lorlatinib-treated cells were poorly dividing, DMSO-treated cells divided normally, which increases the levels of mutations. These findings clearly indicate that drug treatments (a) trigger generation of mutations, (b) patterns of these mutations are treatment-specific, and (c) the mutation patterns are defined not by the chemical structure of the drug, but rather by its target.

### Non-random nature of common mutations

Considering the null hypothesis that identical mutations in independent subclones are random, we calculated the probabilities of their generation. We realize that with such null hypothesis, we may underestimate the probability of “random” drug-induced mutations, since some drugs, e.g. 5FU, show mutation signatures, i.e. cause mutations in a defined nucleotide context. Therefore, potentially, we can overestimate the non-randomness of the mutations. Nevertheless, the limitations of nucleotide context of the mutations can affect only the size of the set of positions that can be mutated (the universal set of cardinality U, see Mathematical analysis in Materials and Methods), which causes a relatively minor effect on the overall calculations of probabilities.

A critical difference between our experiments and conventional evolutionary experiments, where selection was studied, is that all SNPs that we identified were generated *de-novo* in the course of drug treatments, since in calling mutations, we only consider differences between treated and parental clones. In experiments with dose escalation (setup 1), all mutations detected in a subclone must be generated in the previous drug-treatment-recovery cycles. In experiments with a single treatment (setup 2), all mutations must be generated during the time following the application of the treatment and prior to collection of the sample.

A simple example to calculate the probabilities of mutations is based on the experiment with osimertinib, where DNA sequencing was done with the parental and three individual resistant subclones. The resistance was achieved during 9 consequent drug exposures with dose escalation, and the total number of generated mutations in a resistant subclone was 3469 (in subclone 1), while the number of mutations common for all three resistant subclones was 402. Therefore, on average, about 385 mutations should be generated following each treatment cycle in a cell that eventually will give rise to the final subclone (3469 mutations ÷9 treatments, see Table 3), and 45 of them (402 common mutations ÷9 treatments, Table 3) should be identical in three independent subclones. These considerations estimate an average number of mutations per cycle, and it is possible that at some cycles fewer mutations were generated. However, in such a case, higher numbers of mutations were generated at different cycles to account for the overall set of mutations.

We performed a statistical analysis to estimate the probability of random generation of identical mutations, considering their number, the overall number of mutations and the genome size. This probability per cell is extremely low (*p*=7×l0^−468^), see explanation in Mathematical analysis of mutation frequencies, Materials and Methods. Since the total number of cells in each treatment was about 10^6^, the odds of random generation of this number of common mutations are almost non-existent. Similarly, a very strong deviation from the null hypothesis of random mutagenesis was found with doxorubicin (*p*=2×l0^−7792^), since the percentage of recurrent mutations was higher (Fig. 2a). Therefore, these mutations must be attributed to a non-random mechanism(s). To visually illustrate this lack of randomness, Fig. 2g shows a computer simulation of random generation of such mutations. In 100,000 pairs of randomly generated sets of 385 mutations (the average number of mutations generated in one round of osimertinib treatment), the maximum overlap was 1 mutation in a small subset of the pairs, while the experimentally observed overlap was 45 mutations. The probability of generation of this number of common mutations randomly is vanishingly low (*p*=10^−1709^).

### Recurrent mutations are generated during the pseudo-senescence stage

Following administration of the above-mentioned drugs or upon glucose deprivation, we observed similar cell responses, where cells underwent growth arrest and entered a prolonged senescence-like state, followed by the recovery and the resumption of growth. Since this observation suggests that cells become adapted to drugs while being in the pseudo-senescent state, we investigated at what point cells generate the mutations. We collected (a) cloned H1975 cells before the treatment, (b) cells at the pseudo-senescent state on day 4 after addition of osimertinib, and (c) cells at the pseudo-senescent state on day 21 after addition of osimertinib (about 10 days before the beginning of recovery in this experimental setting). Whole genome DNA was isolated and sequenced. Since cells in the treated samples were in pseudo-senescence and therefore did not grow, we sequenced the mixed populations of cells in each sample without subcloning (Fig. 1a, setup 2). We observed that compared to the parental clone there were 4082 SNPs on day 4 and 5858 SNPs on day 21 of pseudo-senescence (Table 4).

Notably, though in this experiment, the cells were not cloned, did not reach the recovery stage, and were exposed to only one treatment cycle, a fraction of mutations overlapped with mutations generated in a subclone of osimertinib-resistant cells in the independent experiment described in table 3 (Fig. 2h). Even higher fraction of the overlapped mutations was seen with mutations common for the three osimertinib-resistant subclones (Fig. 2i). These data further confirm the drug target-specific patterns of mutations, since practically no overlap was seen with mutations generated upon development of resistance to drugs that affect different targets, e.g. osimertinib and doxorubicin (see Fig. 2d,e).

Since cells in the osimertinib treatment day 4 and day 21 samples represented distinct biological replicates that were cultured on separate plates, all these SNPs were generated independently of each other. We observed 803 SNPs that were shared between the two replicates, indicating the recurrency of the mutations. Furthermore, since on average, the sequencing coverage was 30X, our mutation detection limit was about 10% (at least three reads with the mutation out of thirty, Table 2). In other words, to detect a mutation, at least 10% of cells in the population should carry it, which suggests that all detected mutations (even those seen in only one sample) are independently generated in multiple cells in the populations, i.e. they are recurrent. Therefore, recurrent SNPs are generated while cells are in a pseudo-senescent state. Furthermore, they continue gradually accumulating at this non-dividing state in the absence of selection (see section Dynamics of generation of mutations, below).

Similar analysis was performed with cells that developed adaptation to glucose deprivation (glucose deprivation resistance - GDR). In this experiment WGS was performed with parental clone, cells at 10 days in pseudo-senescence and recovered population that resumed growth without glucose. We observed 1002 SNPs common between pseudo-senescent and recovered cells (Table 4). Notably, these SNPs were generated independently, since cells that became pseudo-senescent and that became fully adapted underwent the treatment on different plates. Therefore, as with osimertinib, adaptation to glucose deprivation is associated with active generation of mutations at the pseudo-senescent stage, and a large percentage of these mutations was identical in two independent samples (pseudo-senescent and recovered).

### The mutation recurrency cannot be explained by selection

Since pseudo-senescent cells are neither dividing nor dying, in general, they should not be under significant selection pressure, and therefore the selection is unlikely to explain the recurrency of the mutations. However, we decided to test this directly. We individually barcoded the cells in the parental clone using a Cellecta CloneTracker 50M Lentiviral Barcode Library (Cellecta, inc.), exposed them to glucose deprivation and sequenced barcodes in populations of cells in pseudo-senescence and after the resumption of growth in the glucose-free medium. Barcodes were sorted according to their representation in each population and plotted on a graph. If the cells underwent selection following glucose deprivation, we expected to find a significant overrepresentation of a group of barcodes corresponding to the selected clones in the treated population compared to control (Fig. 3a). Indeed, we see such an effect with cells that recovered from the glucose deprivation and resumed growth on the glucose-free medium (Fig. 3b), i.e. some clones were strongly overrepresented, while others underrepresented. However, the representation of barcodes in pseudo-senescent cells was indistinguishable from that in the control population (Fig. 3b). This result indicates that cells in the pseudo-senescent state did not undergo significant selection (Fig. 3a). Considering that a large fraction of the recurrent mutations was detected at this stage, their recurrency was defined purely by their precise generation without a significant contribution of selection.

**Fig. 3.**
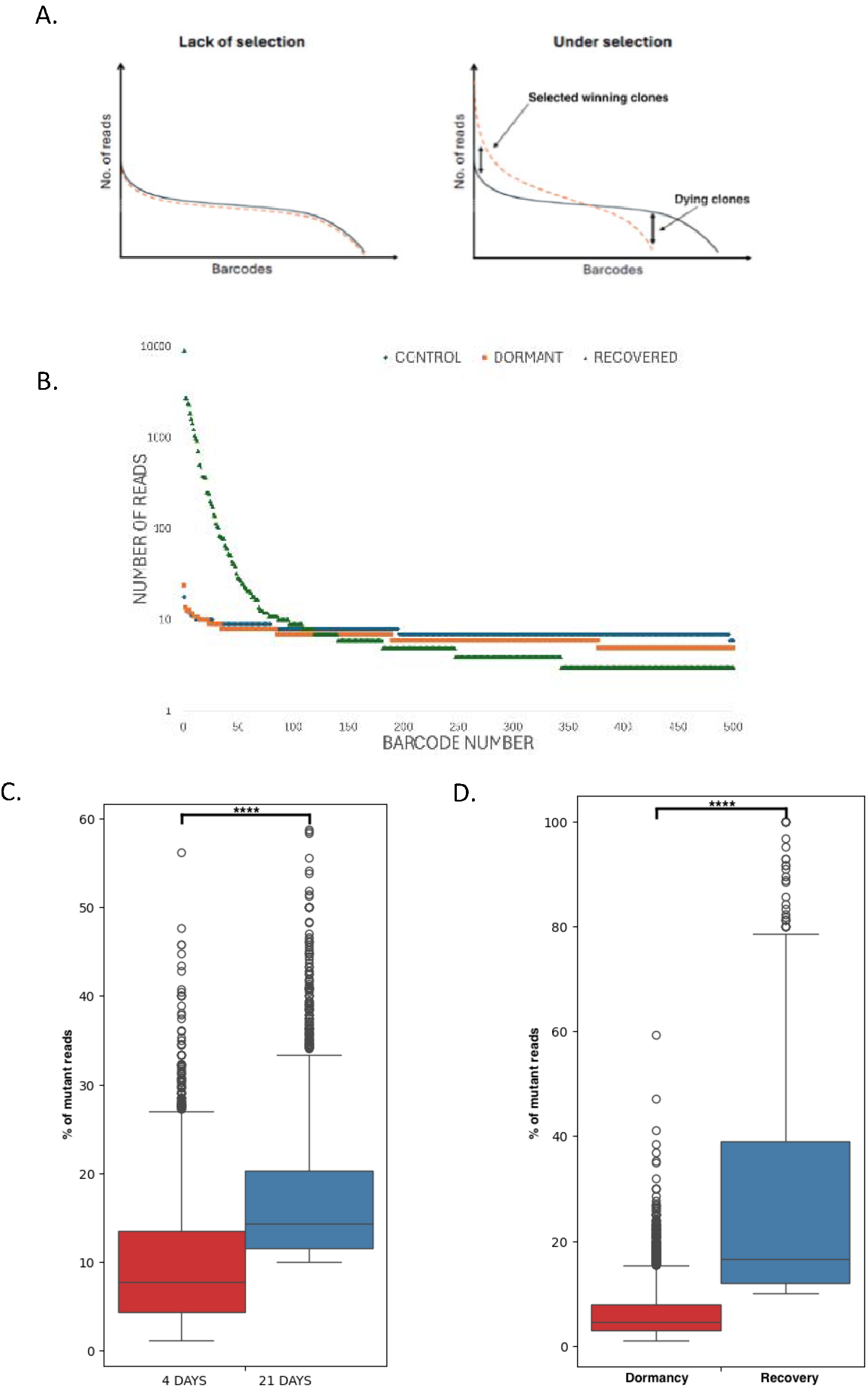
Analysis of mutations in the dormant stage. A. Theoretical changes in the representation of barcodes upon drug or stress treatment without (left panel) and with (right panel) selection. B. Representation of barcodes in dormant and recovered population of cells following glucose deprivation. HCT116 cells were individually barcoded. Following glucose deprivation, cells were collected at the stage of pseudo-senescence at day 10 post-treatment, and after the recovery at day 31. Barcodes were isolated by PCR from the genomic DNA and sequenced by NGS. Representation of the barcodes indicate the lack of enrichment of clones in senescent cell population and strong enrichment of certain clones in the recovered population. C. Representation of reads corresponding to mutations in dormant populations of osimertinib-treated cells at days 4 and 21 of senescence. Left box - frequencies at day 4 time point, right box - frequencies at day 21 time point. D. Representation of reads corresponding to mutations in dormant and recovered populations of GDR cells. Left box - frequencies at day 10 time point (pseudo-senescence), right box - frequencies at day 31 time point (recovery). Unlike mutations in the dormant cells, some mutations in the recovered cells are represented at rates above 40%.

### Dynamics of generation of mutations

Sequencing of mixed cell populations (setup 2) without subcloning allowed us to make estimates of the dynamics of generation of individual mutations over time. Indeed, frequency of a mutation should correspond to the fraction of reads that carry such mutation out of the overall number of reads that cover this specific position in the genome. Accordingly, we analyzed fractions of reads with *de-novo* SNPs at different time points during adaptation to osimertinib and to glucose deprivation. We were focusing on the mutations common for days 4 and 21 of pseudo-senescence following exposure to osimertinib. In this analysis, to account for less frequent common mutations in the early time-point, we reduced the stringency of the mutation filtering at this time point (day 4) by removing the requirement of 10% of the mutant reads. At day 4 pseudo-senescence we observed a broad peak of the fractions of reads corresponding to individual common mutations with median at 8% of reads (Fig. 3c). The median of the fractions of reads corresponding to the same mutations at day 21 was 14% (Fig. 3c). Most significant difference in the read representation was seen with higher frequency mutations (20% and higher). This population of common mutations was dominated by mutations found at 21^st^ day of pseudo-senescence (Fig. 3c). Considering the lack of proliferation (and, subsequently, lack of selection) during the pseudo-senescence stage, this shift indicates continued generation of common mutations. Similar phenomenon was seen with the glucose deprivation experiment (Fig. 3d).

This analysis also revealed an unexpected feature of common mutations to be generated with specific frequencies. If we focus on lower frequency mutations at the 21^st^ day of the osimertinib-treated sample (10-19% of reads, median frequency 13.7%), the median frequency of the corresponding mutations in the day 4 sample was 13.9% (Fig. 4a, 4d). However, if we consider higher frequency mutations in the 21^st^ day osimertinib sample (20-39% of reads, median frequency 26%), the median frequency of the corresponding mutations in the day 4 sample was 18.4% (Fig. 4b, 4d). Finally, upon selection of the highest frequency mutations having 39-60% of reads in the 21^st^ day sample (43% median), the median of the corresponding mutations in day 4 sample was 21.2% (Fig. 4c,4d). Therefore, different recurrent mutations have mutation-specific features of appearing with lower or higher frequencies seen across independent samples and different time points.

**Fig. 4.**
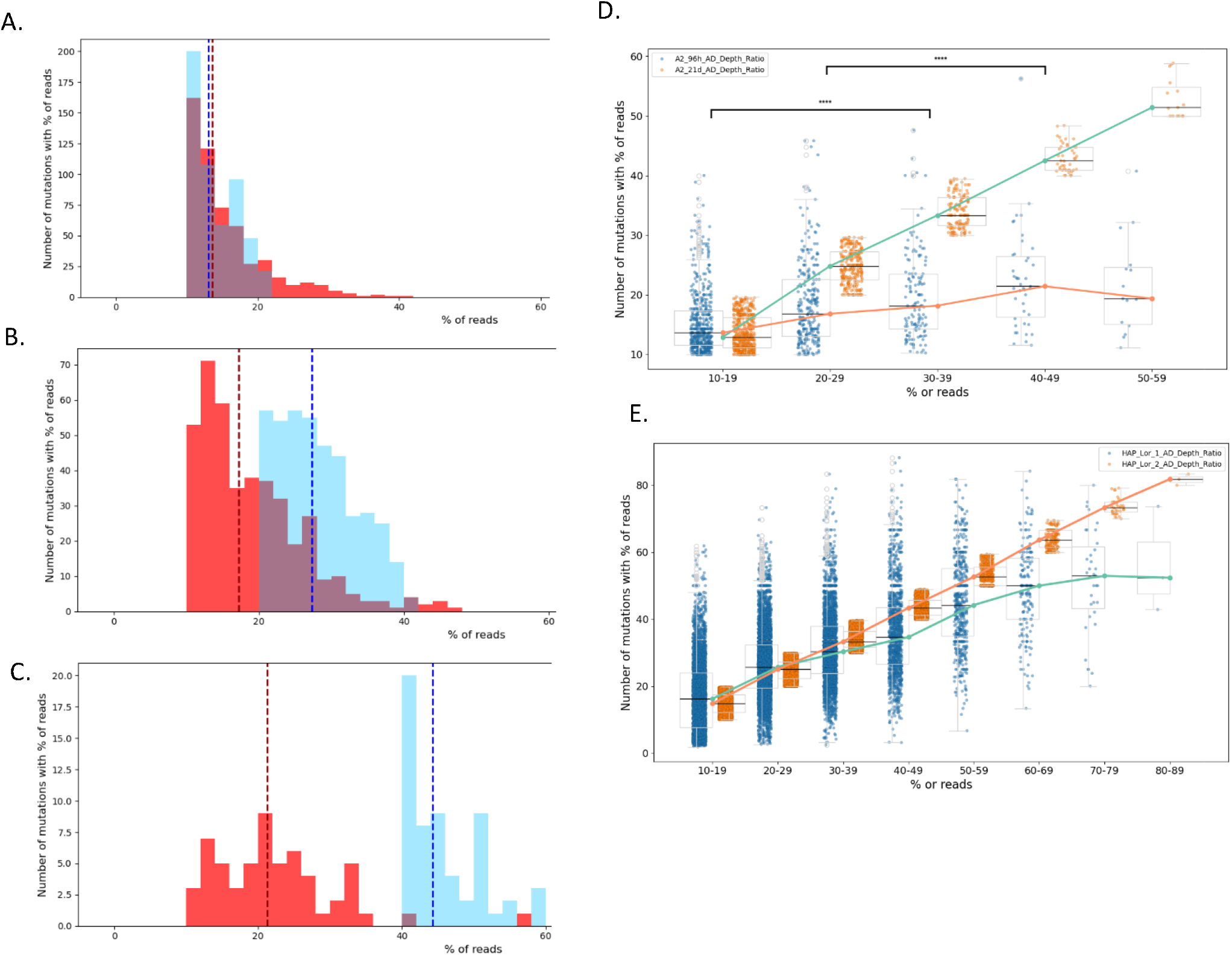
Mutations with different frequencies. A. Analysis of mutations common for days 4 and 21 of senescence. Mutations that show 10-20% representation at day 21, demonstrate lower representation at day 4. Red color - frequencies at day 4 time point, blue color - frequencies at day 21 time point. Grey - overlapping frequencies. B. Mutations with 20-40% representation at day 21 sample and the corresponding mutations at day 4 sample. C. Mutations with 40-60% representation at day 21 sample and the corresponding mutations at day 4 sample. D. Quantification of data presented in Figs. 4a, b, c. Orange-mutations at day 21, blue - corresponding mutations at day 4. Median values for each point are shown as horizontal bars. Statistical difference (paired t-test) between mutation representation in 4-day samples corresponding to 10-20% and 30-40% mutations at day 21 was 4*10^− 5^. Statistical difference (paired t-test) between mutation representation in 4-day samples corresponding to 20-30% and 40-50% mutations at day 21 was 2.9*10^− 3^. E. Analysis of mutations common for sample 1 and sample 2 cells treated with lorlatinib. Sets of mutations with increasing frequencies in samplel and corresponding to them mutations in sample 2. Orange - focused mutations in sample 1, blue - corresponding mutations in sample 2. Median values for each point are shown as horizontal bars.

Similar analysis was performed with two parallel lorlatinib-resistant samples. Unlike osimertininb treatment experiment, where samples were taken at two timepoints in the absence of proliferation, here we compared mutations in the independent recovered populations. If we focus on lower frequency mutations in the lorlatinib sample 1 (10-19% of reads, median frequency 17.3%), the median frequency of the corresponding mutations in sample 2 was 20.2% (Fig. 4e). However, if we consider higher frequency mutations in sample 1 (20-29% of reads, median frequency 26.0%), the median frequency of the corresponding mutations in sample 2 was 27.1% (Fig. 4e). Finally, upon selection of the highest frequency mutations having 70-79% of reads in sample 1 (76.3% median), the median of the corresponding mutations in sample 2 was 52.2% (Fig. 4e). Therefore, it appears that frequencies of generation of mutations at different positions differ significantly, and in some positions can reach almost 100%.

Altogether, these data indicate that mutations accumulate progressively in the pseudo-senescent state before completion of adaptation and resumption of growth, and that their frequencies are orders of magnitude higher than frequencies of spontaneous or induced random mutations or hot spot mutations. Overall, these mutations are not selected at the pseudo-senescent stage, but generated at precise positions with extremely high frequencies, indicating the existence of mechanism(s) of such precise mutation generation.

### Enrichment of mutations associated with changes in DNA methylation

Our next goal was to obtain clues towards possible functional significance of mega-frequency mutations. Therefore, we addressed whether these non-random mutations affected specific functional regions of the genome, and with most treatments, we observed a significant fraction of the mutations in the enhancer areas and thus assessed their association with changes in the DNA methylation patterns. Accordingly, we performed bisulfide sequencing of DNA from cells selected for resistance to osimertinib and glucose deprivation. Comparing cells in pseudo-senescence state and parental clones, we observed no changes in DNA methylation, while cells that achieved osimertinib resistance had 50,000 differentially methylated sites and GDR cells had 75,000. Among the differentially methylated sites, 25.2% for osimertinib and 26.9% for GDR were mapped to enhancers. Therefore, we assessed whether these enhancers preferentially carry the mutations. Indeed, a significant enrichment of mutations in enhancers that carried differentially methylated cytosines was seen with both osimertinib-resistant and GDR cells (Table 5). The enrichment was seen with both hyper and hypomethylated sites (Table S3, S4). This enrichment was seen in active enhancers previously identified (in ENCODE) in HCT116 cells, which we used in the glucose deprivation experiment (Table S5).

**Table 5.** Correspondence of mutations to methylation patterns in GDR and Osi treated cells.

|  |  |  |
| --- | --- | --- |
| Number of Enhancers | 345341 | 345341 |
| Differentially methylated Enhancers | 12197 | 8703 |
| **% of Diff. methylated Enhancers** | **3.4** | **2.5** |
| Mutations in Enhancers | 4768 | 377 |
| Mutations in Diff. methylated Enhancers | 467 | 36 |
| **% of Mutations in Diff. methylated Enhancers** | **9.8** | **9.5** |
| **Fisher's Exact test (p-value)** | **1.04E-88** | **1.1E-11** |

The association of the mutations with enhancers that carry differentially methylated cytosines suggests that either DNA methylation could influence generation of the mutations, or the mutations can influence changes in the methylation patterns. The fact that in pseudo-senescent cells there were no changes in DNA methylation, while mutations were already present at this stage, suggests that the changes in DNA methylation are secondary to the generation of mutations, and therefore mutations may influence the methylation dynamics in the enhancer regions.

### Association of the mutations with binding sites of certain transcription factors

To further assess potential biological effects of the mutations, we assessed their association with binding motifs of transcription factors using JASPAR database. The q-value threshold for the association was set at 0.05. In this analysis, since the transcription factor binding motif can be 15bp and even longer, we established a window of 32 bp at each side of the mutation and identified the binding motifs located within this window. To identify the enrichment of association of the mutations with the transcription factors binding motifs, we randomly distributed the 65bp mutation windows (32bp-1bp SNP-32bp) over the genome in silico. The sizes of the random mutation windows sets corresponded to the numbers of the experimental mutation windows in our datasets. We further calculated whether association of mutations in our experimental datasets with the binding motifs is enriched compared to randomly distributed mutation windows. This test clearly demonstrated a highly significant enrichment of mutations located either within or near transcription factors binding motifs. This association was seen with various drug-resistant cells (Fig. 5a).

**Fig. 5.**
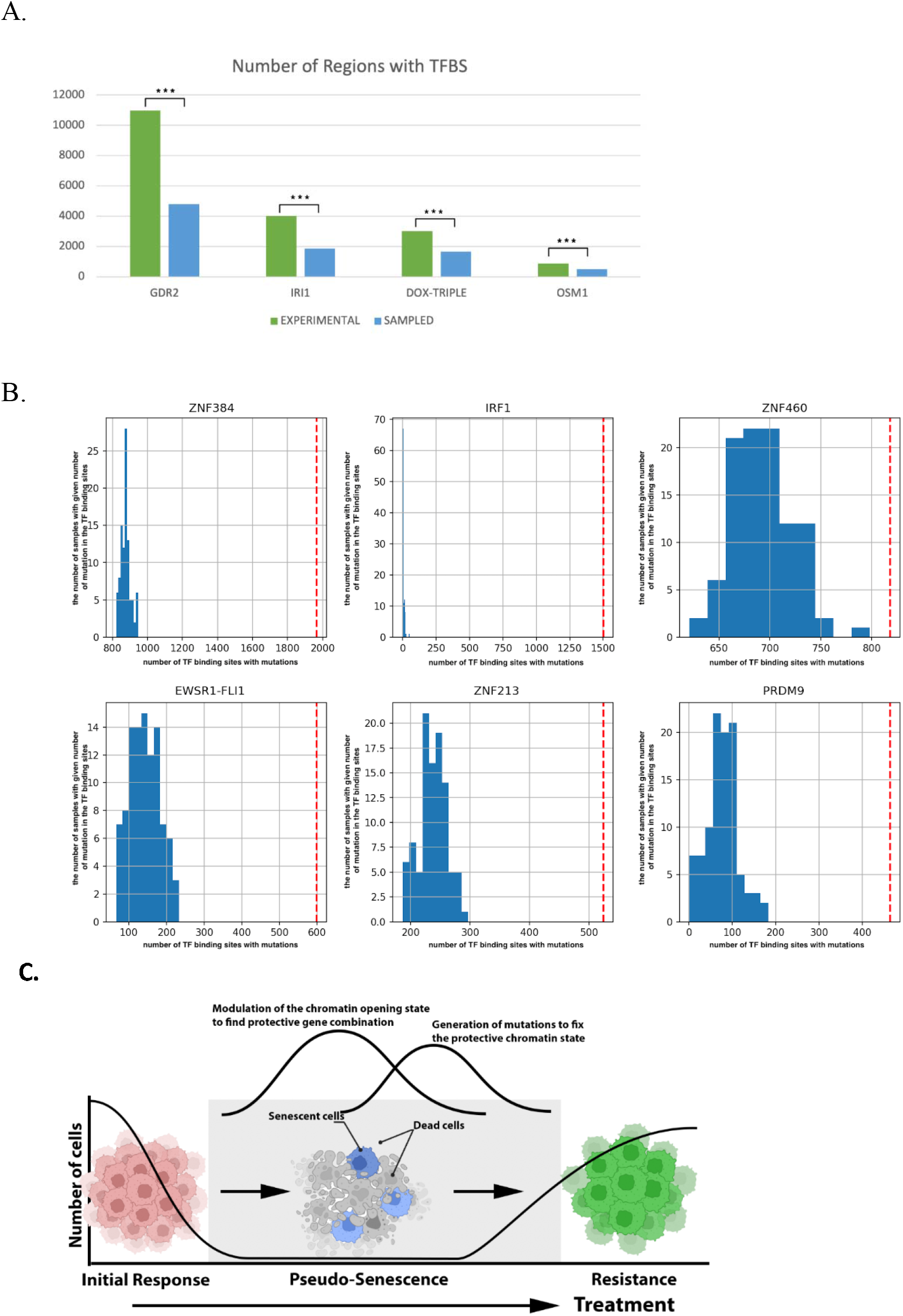
Enrichment of mutations within and near the transcription factors binding motifs. A. Enrichment of mutations within and near the total number of transcription factors binding motifs. Random *in silica* sampling of the 65bp mutation window along the genome served as control. Significant binding motifs were determined according to JASPAR database. The following mutation sets are shown: GDR2 - GDR recovery point, IRI1 - irinotecan subclone 1, DOX-TRIPLE - Doxorubicin common mutations in three subclones, 0SM1 - osimertinib subclone 1. B. Enrichment of mutations within and near binding motifs of certain transcription factors. Here, we show the data for Lorlatinib. Data for other treatments is presented in Table 6. Observed numbers of mutations associated with the binding motif of a transcription factor are shown as red lines. Random *in silica* sampling of the 65bp mutation window along the genome served as control (100 samples) - blue columns. X-axis - number of binding sites. Y-axis - number of random samples with the shown number of binding sites. C. Model of the role of generation of mutations in adaptation to drugs. In pseudo-senescent stage cells modulate their chromatin opening state to find a gene combination that can allow the treatment survival. This state of the chromatin is then “fixed” by mutations in the regulatory regions.

Further, we tested which transcription factor binding motifs associate with the mutations. In this analysis, we also used random sampling of the 65bp mutation window along the genome; the number of windows in each sampling corresponded to the number of mutations in each drug condition; and the number of random samples was 100. This analysis demonstrated that mutations in our datasets were highly associated with several transcription factors binding motifs (Table 6, Fig. 5b). There were common highly enriched motifs seen with all drug-resistant cells, e.g. IRF1, RREB1, KLF9. Notably, almost 75% of IRF1-associated mutations were located strictly within the binding motifs. In contrast, only 28% of KLF9-associated mutations were located within the binding motifs. RREB1 showed intermediate fractions of mutations within the binding motifs. Interestingly, other motifs were enriched only in cells resistant to specific drugs, e.g. SP1 or KLF1 with osimertinib-resistant cells, or FOXD3 and PRDM9 with lorlatinib-resistant cells. Therefore, with different drug and stress treatments, precise mutations are generated preferentially in or near binding motifs of certain transcription factors that may significantly contribute to the transcriptome changes and facilitate drug adaptation.

**Table 6.** Enrichment of mutations within and near the transcription factors binding motifs. Red dots correspond to high significance p<0.01, blue dots - p<0.05. OSM 2TP-osimertinib, second time point - day 21 of senescence. OSM subclone - osimertinib, subclone 2.

| TFBS | OSM<br>2 TP | LOR | CRIZ | GDR | OSM<br>SUBCLONE | IRI | DOX |
| --- | --- | --- | --- | --- | --- | --- | --- |
| RREB1 | • | • | • | • | • | • | • |
| ZNF384 | • | • | • | • | • | • | • |
| IRF1 | • | • | • | • | • | • | • |
| ZNF213 | • | • | • | • |  |  | • |
| ZNF135 | • | • | • | • |  | • | • |
| ZNF460 | • | • | • | • |  | • | • |
| KLF9 | • |  |  | • | • | • | • |
| ZNF740 | • |  |  | • |  |  |  |
| PATZ1 | • | • | • |  |  |  |  |
| ZNF281 | • | • | • |  |  |  |  |
| PRDM9 | • | • |  |  |  |  |  |
| KLF5 | • |  |  |  |  |  |  |
| ZNF148 | • |  |  |  |  |  |  |
| KLF10 | • |  |  |  |  |  |  |
| SP1 | • |  |  |  |  |  |  |
| SP4 | • |  |  |  |  |  |  |
| KLF14 | • |  |  |  |  |  |  |
| KLF12 | • |  |  |  |  |  |  |
| SP2 | • |  |  |  |  |  |  |
| ZNF320 | • |  |  |  |  |  |  |
| ZSCAN4 |  |  |  | • |  | • | • |
| ZNF93 |  |  |  | • |  |  |  |
| EWSR1-FLI1 |  | • | • |  |  |  |  |
| SPI1 |  | • |  |  |  |  |  |

## Discussion

According to classical understanding of mutations in cancer, they arise randomly (41). In human tumors, genomic instability, MMR/HR deficiency, and oncogenic signaling activation increase the level of mutagenicity, thus providing a broad spectrum of variants from which clones with increased resistance to the changed microenvironment/therapy can be selected (42,43). Targeted therapy (BRAF, EGFR, ALK, MEK inhibitors, etc.) often results in secondary mutations in the drug target itself or in downstream components of the pathway. These mutations were considered to accumulate either by selection of preexisting subclones or by *de novo* generation of random mutations during the treatment, similarly, followed by selection (41,42,44).

Here, we report that upon adaptation to drugs, non-growing cancer cells actively generate multiple non-random mutations at very specific locations and that the patterns of these mutations are specific to the drug treatment or stress. Notably, the phenomenon of generation of recurrent drug-induced mutations is fundamentally different from the known phenomenon of hot spot mutations, because the latter usually occur with frequencies 10-100 times higher than average (i.e. 10 ^−7^ − 10^−8^, rather than normal 10^−9^), while frequencies of these drug-induced mutations could be several orders of magnitude higher (i.e. 10^−1^ and higher), see also section Dynamics of generation of mutations.

This striking and unusual generation of non-random mutations with extremely high frequencies raises the question why previous cancer genome sequencing studies overlooked this phenomenon? Probably, the main reason is that previous studies were directed towards the discovery of unique mutations affecting protein structure or function that confer drug resistance, e.g. resistance to osimertinib or lorlatinib (45-47). This led to even stronger focus on mutations in the drug targets, e.g. EGFR or ALK, or on genes that belong to pathways that were known from previous studies to cause drug resistance (48-50). Accordingly, exome sequencing, rather than whole genome sequencing, was performed in most of these studies, which automatically eliminated all mutations that arise in non-coding regions, including regulatory sequences, like enhancers. Overall, in these genetic analyses the goal was not to investigate genetic changes that take place during drug adaptation in an unbiased way, but to identify functionally important mutations, which usually required prior knowledge about the role of specific genes or pathways. In these studies, “orphan” mutations without clear functional roles were simply disregarded as unimportant. In contrast, our approach was to identify all SNPs in the genome in an unbiased way, which led to the discovery of a novel phenomenon – generation of precise drug-specific mutations.

Why mutations with frequencies that can reach over 50%, many orders of magnitude higher than hot spot mutations, were overlooked in previous studies? A possible reason is that previously, investigators were not dealing with a system that lacks selection. Accordingly, high frequency mutations were considered to result not from a high frequency of their generation, but rather from selection. In addition, the calculation of the frequencies is usually done by looking at the number of mutations per generation normalized by the size of the genome. We, on the other hand, calculate the frequencies of the generation of mutations based on the fraction of cells that acquired these mutations. This approach is valid only because in our system, there is no cell propagation and selection.

An important observation with two different Alk inhibitors crizotinib and lorlatinib was that the patterns of mutations were relatively similar, indicating that the patterns are defined by the nature of the drug target, rather than by the chemical structure of the drugs. This finding strongly suggests that the generation of specific mutations results from the downstream signaling events of the inhibition of the drug target and is unrelated to potential direct chemical effects of the inhibitory molecules on the DNA repair systems.

Another important finding was that a significant fraction of the mutations is generated when cells are staying in the pseudo-senescent state prior to adaptation and resumption of growth. During this stage, cells are modulating gene expression patterns that can allow them to survive and start dividing. The fact that the mutations are generated during this period suggest that they may significantly contribute to survival.

The main questions arising from this study are (a) what is the mechanism of generation of these drug-specific mutations, and (b) what is their role in drug adaptation. To address possible mechanisms of generation of the recurrent mutations, we performed a COSMIC mutation signature analysis. The most prominent and common to different treatments signature was SBS5. This is a very common signature in cancer samples with unclear mechanism of generation (55). Therefore, this approach did not provide a clear answer. We propose that there must be some connection between the precise generation of mutations and the chromatin structure. We can rule out a connection to DNA methylation, since the majority of mutations were not physically linked to positions of C that underwent methylation or demethylation. Therefore, we suggest that there may be some other event, probably a kind of histone modification that can precisely target mutations to specific positions. Another possibility is the link to binding of a transcription factor. For example, it was reported that C/EBP can facilitate recruitment of APOBEC to specific positions (56). Possibly, similar mechanisms may function in generation of our precise mutations, though obviously in addition to APOBEC, generation of our mutations must be governed by other mechanisms, possibly components of DNA repair pathways.

A reasonable possibility regarding the function of the mutations is that they may affect the state of the chromatin to facilitate adaptation to drug/stress conditions. A hint about possible chromatin changes associated with the mutations came from the analysis of changes of global DNA methylation upon development of resistance to glucose deprivation and osimertinib. Indeed, upon comparison of differential CpG methylation events and the presence of the mutations in enhancers, there was a significant enrichment of differentially methylated sites in the mutated regions. These data suggest that the mutations may affect the CpG methylation status of the regulatory regions.

Another important observation was that the mutations were highly enriched within binding motifs of certain transcription factors or in close proximity to them. The mutations may influence expression of genes regulated by these transcription factors. Furthermore, they may affect the chromatin state in these regions as secondary effects via binding of these transcription factors, since some of them, e.g. SPI1, FOXD3 or PRDM9, can function as pioneer transcription factors in recruiting chromatin remodeling factors (51-54).

Notably, since the number of drug-induced mutations and downstream genes is very high, each of the mutations may have a very small effect on gene expression, which could be below our detection limits. However, together they may produce cumulative biologically significant effects on providing the resistance.

Overall, we demonstrated that (a) multiple mutations are actively generated upon stress or drug exposure, (b) a large fraction of them is recurrent, (c) mutation patterns are drug-target-specific, indicating the role of downstream physiological responses in their generation, (d) accumulation of mutations takes place at pseudo-senescence stage, when cells develop the resistance, (e) accumulation of mutations does not involve selection, (f) mutations are enriched in binding sites of certain transcription factors and associate with changes in the methylation states of enhancers. We propose that upon drug adaptation, when cells experience extreme stress, some changes in chromatin may trigger local specific mutations by recruiting APOBEC family members or causing local DNA damage. Such generation of mutations preferentially takes place in the proximity to or within binding sites of certain transcription factors, which may either increase or decrease their activity, leading to multiple changes in transcriptome that may facilitate the adaptation.

Our major hypothesis is that the recurrent mutations may provide an additional layer in drug protection making it permanent. Indeed, in line with previously discussed models, following drug treatment in a dormant or pseudo-senescent stage, cells modulate chromatin state and expression of various genes attempting to find a combination that provides the ability both to survive this drug concentration and to resume growth. We hypothesize that when such a protective chromatin state is achieved, cells generate mutations in the regulatory regions that genetically fix this chromatin state and gene expression pattern (Fig. 5c) and thus make the protection permanent.

## Materials and Methods

### Cell culture and reagents

HCT116 cells (ATCC Cat# CCL-247, RRID:CVCL_0291) were obtained from the American Type Culture Collection and maintained in McCoy’s 5A medium supplemented with 10% fetal bovine serum (FBS), 4 mM L-glutamine (BI-Biologicals, Cat#03-020-1B), 2 mM L-alanyl-L-glutamine (BI-Biologicals, Cat#03-022-1B), and 1% penicillin-streptomycin (BI-Biologicals, Cat#03-031-1B). H1975 cells (NCI-H1975) were obtained from the American Type Culture Collection and maintained in RPMI-1640 medium supplemented with 10% FBS, 4 mM L-glutamine, 2 mM L-alanyl-L-glutamine, and 1% penicillin-streptomycin. NSCLC cell line H3122 was obtained from the Lung Cancer Center of Excellence Cell Line depository at Moffitt Cancer Center. H3122 cells were grown in RPMI-1640 containing L-Glutamine (Gibco, Cat#25030081), supplemented with 10% fetal bovine serum (FBS), 1% penicillin-streptomycin, and 10μg/ml human recombinant insulin (Gibco, Cat#12585014). Cells were incubated at 37°C with 5% CO_2_ atmosphere with constant humidity. Cells were passaged as needed using 0.05% trypsin (Gibco Cat#25300054).

Conventional therapy drugs were purchased from Sigma-Aldrich (St. Louis, MO, USA): Doxorubicin (25316-40-9) and SN-38 (<u>86639-52-3</u>). Targeted therapy drugs were purchased from MedChem (Monmouth Junction, NJ, USA): osimertinib (HY-15772 ), Crizotinib (<u>HY-50878</u>) and Lorlatinib (HY-12215).

### Selection of the resistant clones

ALK inhibitors: 2 million of H3122 cells were plated on 10cm plate. Naïve cells were treated with 0.5uM crizotinib, 2.0uM lorlatinib, or 0.025% vehicle (DMSO) (Sigma, Car#D8418) for four weeks with media and drug or drug vehicle changed every 3-4 days. Cells in drug vehicle were passaged as needed during the length of the experiment. After four weeks, cells under drug had multiplied and started to cover the plate, indicating resistance to the drugs.

Osimertinib: 1 million of H1975 cells were plated on 10cm plate and 6mM osimertinib was added for 24h. This led to the death of about 50% of the population and pseudo-senescence of the rest of the population with complete growth cessation. In about three weeks, cells started to multiply and cover the plate. The treatment was repeated either with the same or higher doses of osimertinib. After nine cycles of treatment with dose escalation, we obtained cell that could grow after the treatment with 18mM of the drug.

Similarly, using multiple cycles of treatment with dose escalation, we selected HCT116 cells that could grow after the 24h treatment with 40 nM irinotecan or 500nM doxorubicin.

Glucose deprivation: 1 million of HCT116 cells were plated on 10cm plate and the medium was replaced to no-glucose medium. For about four weeks the media was changed every 3-4 days. After four weeks, cells started to multiply and cover the plate, indicating that GDR cells took over the population.

### Cell cloning

To establish monoclonal populations, cells were subjected to limiting dilution cloning. Individual clones were selected using cloning discs (Merck, Cat# Z374431), then expanded and cryopreserved in liquid nitrogen in a freezing solution composed of 10% dimethyl sulfoxide (DMSO) in fetal bovine serum (FBS) for long-term storage.

### β-gal activity assay

β-gal activity in the senescence cells were observed using the Senescence Beta-Galactosidase staining kit (Cell Signaling Technology, Cat#9860S) according to the protocol manual.

### DNA barcoding

Barcoding of the cloned cell population was performed using the Cellecta CloneTracker 50M Lentiviral Barcode Library (RRID:SCR_021827), in accordance with the manufacturer’s protocol. Cells were transduced with the lentiviral barcoding library and subjected to puromycin selection to enrich for successfully infected cells. Following selection, cells were allocated into separate groups for drug treatment. After recovery, genomic DNA was extracted, and the integrated barcodes were amplified via nested PCR (38). All resulting samples were pooled and prepared for high-throughput sequencing.

### DNA extraction and library barcodes analysis

Genomic DNA was extracted from cultured cells using the Wizard Genomic DNA Purification Kit (Promega, Cat#A1120). Barcode sequences were amplified via nested PCR (38). The first PCR (PCR1) utilized Titanium Taq DNA Polymerase (Takara Bio, Cat# 639209), and PCR products were purified using the QIAquick PCR & Gel Cleanup Kit (Qiagen, Cat#28506). A second round of amplification (PCR2) was performed with nested primers—either generic or sample-specific— using the Phusion High-Fidelity PCR Master Mix (Thermo Scientific, Cat#F531S). A secondary barcode was added during PCR2 to enable sample multiplexing. PCR products were normalized individually, pooled, and further purified using AmpureXP magnetic beads (Beckman Coulter, Cat#A63881) according to the manufacturer’s instructions. Next, the barcodes were sequenced using Ion Torrent.

### Whole-Genome Sequencing and analysis

Genomic DNA samples underwent whole-genome sequencing at 30× average coverage by BGI, Inc., employing next-generation sequencing technology to generate paired-end reads. Raw FASTQ files were subjected to quality control using FastQC (v0.11.7), evaluating per-base quality scores, adapter contamination, and sequence duplication levels to ensure data integrity prior to processing (57).

### Read Alignment and Preprocessing

High-quality reads were aligned to the human reference genome (GRCh38/hg38) using BWA-MEM (v0.7.17) with default parameters, which optimizes for local alignments and handles indel rescue (58). Resulting SAM files were sorted, converted to coordinate-compressed BAM format, and indexed via samtools (v1.16). Local realignment around indels and base quality score recalibration (BQSR) followed GATK best practices to mitigate alignment artifacts and systematic sequencing errors (59).

### GATK Processing Pipeline

Processed BAM files underwent standardized GATK (v4.6.1.0; HTSJDK v4.1.3, Picard v3.3.0) preprocessing, including read group assignment (AddOrReplaceReadGroups), PCR duplicate marking (MarkDuplicates), and covariate-based BQSR (BaseRecalibrator followed by ApplyBQSR). These steps enhance variant calling accuracy by correcting for covariates such as machine cycle, nucleotide context, and read position bias.

### Variant Calling

Multisample variant detection employed GATK Mutect2 in tumor-normal mode, integrating one parental cancer control and treated tumor samples with the following key parameters: germline resource (gnomAD exomes), panel-of-normals (PON), genotyping of germline/PON sites, relaxed LOD thresholds (normal LOD=0, initial tumor LOD=0.1, emit LOD=1), zero minimum allele fraction, and low allele frequency resource prior (1e-6). The mentioned LOD thresholds were set up to obtain maximal number of mutations without a significant number of multiallelic and weak positions. High thresholds in further filtration corrected for possible errors at this stage. Output comprised a multisample VCF with associated F1R2 tar.gz files for contamination and bias modeling.

### Variant Pre-Filtering

Following variant calling, an initial filtering step was applied including artifact modeling (LearnReadOrientationModel), filtering with FilterMutectCalls according to GATK best practices. These steps serve to flag likely false positives while retaining candidate variants for further analysis. It is important to emphasize that these filters are advisory rather than definitive thresholds. Subsequent filtering was performed with criteria tailored to the specific experimental design. For example, in experiments involving populations in a sequence of time points, we retained low-frequency variants in the early time point and validated their presence in subsequent time point to confirm the authenticity. Thus, downstream filtering strategies were customized to preserve relevant variants based on experiment-specific considerations rather than relying solely on generic filter parameters.

### Variant filtering

Filtering was performed using bcftools (v1.21) in the Linux environment. For the mutation filtering, we required at least 20 reads in control sample per position and fully homozygous state in the parental clones. In its standard setting, the software first eliminates low-quality reads and then identifies reads with new alleles that are absent in the parental clone. The problem with this setting is that if the “new” allele is present in some of the low-quality reads in the parental clone, it will be recognized as *de-novo* mutation in the resistant subclone, though it could preexist in the parental clone. Accordingly, for the *de-novo* mutations, we analyzed BAM file and applied a stringent filter by selecting alleles that were not present even in one read in the parental clone, including low quality reads. On the other hand, we applied a requirement for the presence of the mutant allele in more than 20% of reads in resistant subclones. For experiments with mix cell populations without cloning after the treatment, we required (a) 10% of reads and (b) at least three reads in resistant subclones. For the filtered VCF files used in this study, see Tables S6-S21.

### Whole Genome Bisulfite Sequencing and Analysis

The whole genome from cultured cells were isolated by Quick-DNA™ Miniprep Plus Kit (Zymo Cat#D4068) following the manufacturer protocol. Isolated DNA was sent for Whole Genome Bisulfite sequencing as a service to BGI, Hong Kong. Sequencing data were processed using the nf-core/methylseq v3.0.0 pipeline (60). Initial and post-trimming quality were assessed using FastQC. Adapter trimming and low-quality bases were removed using Trim Galore!, which integrates Cutadapt for adapter detection and trimming (61). Trimmed reads were aligned to the reference genome using bwa-meth, a bisulfite-aware wrapper around BWA-MEM (62). Duplicate read marking and associated alignment processing were performed using Picard tools (63). Alignment quality metrics were generated using Qualimap (64). Library complexity and yield projections were assessed using Preseq (65). Finally, all quality control reports were aggregated into a unified summary using MultiQC (66).

### Differential Methylation analysis

Differential Methylation analysis was performed using the methylKit R package (v1.30.0) (67). Raw coverage files for methylation calls at CpG sites for each sample were imported with methRead and further filtered ensure high confidence methylation calls. Filtration was done based on the coverage, where the sites with below 10 reads and more than 99.9^th^ percentile of coverage were discarded for each sample. Following filtration, the normalization of coverage between the samples was performed by using the median method to reduce bias. The samples were then merged to generate a consensus CpG matrix to extract the sites that are covered by reads in all the samples. Next, the samples were further filtered to retain only the sites with standard deviation >2%. Differential methylation between groups was performed using the overdispersed method with the Fisher’s exact test and Benjamini-Hochberg (BH) FDR adjustment for statistical significance. CpG sites were called as differentially methylated if they have difference ≥25% and FDR-adjusted q-value ≤0.05. For the list of differentially methylated sites in osimertinib-resistant and GDR cells compared to control clones, see Tables S22-S23.

### ATAC sequencing and analysis

Samples for ATAC sequencing (ATACseq) were prepared using Zymo-Seq ATAC Library Kit (Cat#D5458) following the manufacturer protocol. The prepared libraries were sent for paired-end sequencing as a service to BGI, China.

Raw ATACseq reads were first check for quality using FastQC (v0.11.7) and trimmed using Trim Galore (v0.5.0). The reads were then aligned to reference genome hg38 with BWA-MEM (v0.7.17) with paird-end default settings. Once the SAMfiles were converted to BAM files using samtools (v1.16), the aligned reads were filtered to remove reads mapping to mitochondrial DNA and blacklist regions, unmapped, non-primary, supplementary and duplicate reads using samtools and picard tools (68,63). The filtered BAM files were used to call peaks using MACS2 (v 2.1.1.20160309) with appropriate parameters for paired end reads (69). For the list of ATACseq peaks in parental HCT116 clone, see Table S24.

### Analysis of mutations in transcription factor binding sites

To assess whether mutations are enriched within transcription factor binding sites (TFBSs) and to identify genomic regions where such enrichment occurs beyond random expectation, we performed the following analysis.

#### Generation of genomic windows around mutations

Mutations were extracted from VCF files and converted into BED format by defining fixed-width genomic windows centered on each mutation. Each window spanned 65 nucleotides, comprising 32 nucleotides upstream and 32 nucleotides downstream of the mutation position. To construct a null distribution, random genomic regions were generated by uniformly sampling genomic positions across the reference genome. For each VCF file, the number of randomly sampled positions matched the number of observed mutations, and the same window size (65 nucleotides) was applied. This random sampling procedure was repeated 100 times to account for stochastic variability.

#### Motif scanning and filtering

Motif occurrence analysis was performed on both observed and randomly sampled genomic windows using FIMO from the MEME Suite (70). FIMO was run with default parameters to identify transcription factor binding motifs within each genomic window. To control for spurious motif matches, only motif occurrences with a q-value ≤ 0.05 were retained for downstream analysis.

#### Quantification of motif occurrences

For each transcription factor motif, the number of unique genomic windows containing at least one occurrence of the motif was counted. Multiple occurrences of the same motif within a single window were counted only once, ensuring that each window contributed at most one count per motif. This procedure was applied identically to the observed mutation-centered windows and to each of the 100 randomly sampled datasets.

### Data Visualization and Statistical Analysis

Data visualization was performed in Python using the pandas, matplotlib and seaborn packages. Statistical analyses were calculated either in the R or python programming language. Fisher’s Exact test or Chi-Squared test were used to evaluate the statistical significance. A P-value of less than 0.05 was considered significant.

For each motif, the distribution of motif counts obtained from random sampling was compared with the experimentally observed count. The observed enrichment was quantified using fold change relative to the mean of the null distribution and by computing a Z-score. Statistical significance was assessed using an empirical permutation-based test. This approach provides a conservative estimate of statistical significance that does not rely on parametric assumptions about the null distribution.

### Datasets Used

For Enhancer analysis the enhancer datasets and enhancers for HCT116 cell line (ENCODE) from GeneCards database were used (71). The Transcription factor binding motifs from JASPAR database were used for analysis of mutations in transcription factor binding sites (72).

### Mathematical analysis of the mutations’frequencies

#### Input

- The size of the human genome - the universal set *U* of cardinality |*U*| *= n=* 3*10^9^.
- The number of mutations in two independent sets of resistant cells (two subclones or two plates with mixed populations) *A* and *B* of cardinality. We assume that |*A*| = |*B*| = *m*

#### Output

- The number of common mutations k equal to intersection between |*A*| and |*B*|. The probability P(|*A* ∩ *B*|= *k*).

Main assumptions

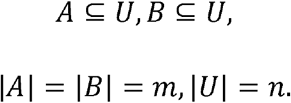

The subsets *A* and *B* are chosen uniformly at random from all subsets of *U* of size *m*. The choices of *A* and *B* are independent.

First exact formulae

We need to compute:

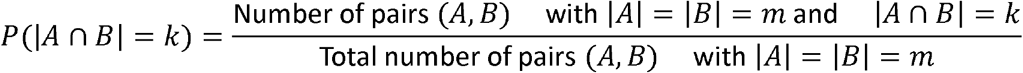

1. The total number of pairs (*A,B*). Evidently, the total number of pairs is

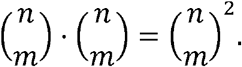
2. The number of favorable pairs. Choose *S* of size *k* for the intersection. Choose *A* to be *S* ∪ *Ã*, where *Ã* is chosen from *U* − *S* of size *m* − *k*. Choose *B* to be 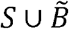, where 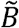 is chosen from *U* − *S* of size *m* − *k*, but we require that 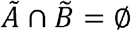 to ensure that the intersection is exactly S.

Hence, the number of ways to choose *S* is 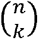.The number of ways to choose *Ã* is 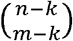.

To determine *B* it is needed to choose 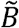 from *U* − *S* of size *m* − *k*, but disjoint from *Ã*.

The number of ways to choose 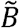 disjoint from *Ã* is

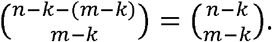

Therefore, the number of favorable pairs is

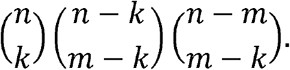

So, the probability needed is

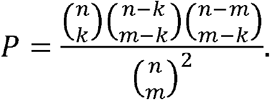

To simplify this expression, note that

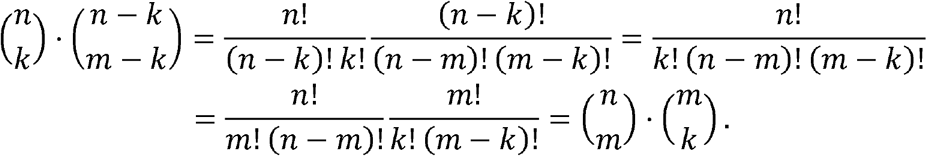

Consequently, the probability is

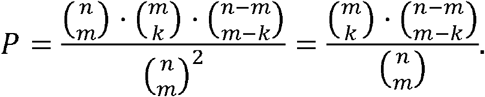

In summary, the probability that two randomly chosen subsets of size *m* from a universal set of size n have an intersection of size *k* is

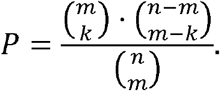

Validity Requirements:

1. 0 ≤ *k* ≤ *m* ≤ *n*,
2. *m* − *k* ≤ *m* − *n*, otherwise 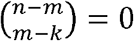

Second exact formulae

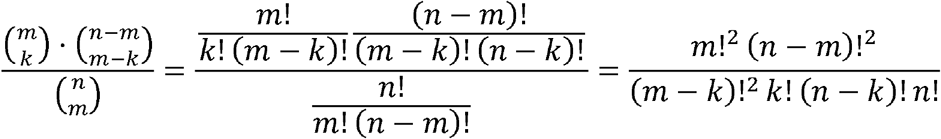

Some known symbolic approximations

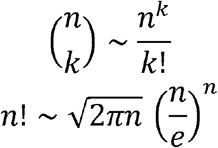

Numerical approximations:

- First calculation - for mutations present in two out of three subclones

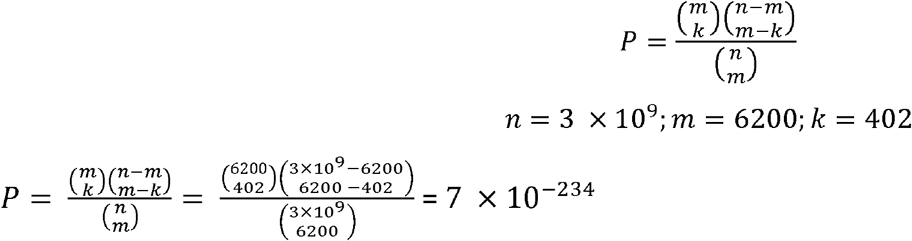
- Second calculation - for mutations present in all three subclones:

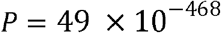

Therefore, in the described process examining mutations following exposure to a drug, the observed patterns are not random.

## Data Availability Statement

All data generated in this work are currently being uploaded and will soon be available in the GEO database.

