## Supplementary material for "Mega-frequency mutagenesis: generation of non-random precise mutations with extremely high frequency upon adaptation of cancer cells to drugs and stress": Table S2, Table S3, Table S4, Table S5

**Supplement Materials**

**Table S1 is an excel file**

| **Table S2** | |  | |  |
| --- | --- | --- | --- | --- |
| Number of ATACseq peaks | | 132000 | |  |
| bp covered by the peaks | | 35540706 | |  |
| Number of mutations | | 34744 | |  |
| Peaks that carry mutations | | 362 | |  |
| bp covered by the peaks with mutations | | 154411 | |  |
| Genome | | 3E+09 | |  |
| **Expected mutations within peaks** | | **411** | |  |
| **Observed** | | **368** | |  |
| **Probability that a mutation falls in a peak randomly** | | **0.0118469** | |  |
| **Table S3** | | **GDR** | | **Osi** |
| Number of Enhancers | 345341 | | 345341 | |
| Hypermethylated Enhancers | 7477 | | 6162 | |
| **% of Hypermethylated Enhancers** | **2.1** | | **1.8** | |
| Mutations in Enhancers | 4768 | | 377 | |
| Mutations in Diff. methylated Enhancers | 331 | | 22 | |
| **% of Mutations in hypermethylated Enhancers** | **6.9** | | **5.8** | |
| **Fisher's Exact test (p-value)** | **8.09E-75** | | **1.7E-06** | |
| **Table S4**  Number of Enhancers | **GDR**  345341 | | **Osi**  345341 | |
| Hypomethylated Enhancers | 4720 | | 2541 | |
| **% of Hypomethylated Enhancers** | **1.3** | | **0.7** | |
| Mutations in Enhancers | 4768 | | 377 | |
| Mutations in Hypomethylated Enhancers | 166 | | 15 | |
| **% of Mutations in hypermethylated Enhancers** | **3.5** | | **4.0** | |
| **Fisher's Exact test (p-value)** | **1.50E-26** | | **2.0E-07** | |

| **Table S5** | **Total** | **Hypermeth.** | **Hypometh.** |
| --- | --- | --- | --- |
| Number of Enhancers | 58561 | 58561 | 58561 |
| Diff.methylated Enhancers | 5603 | 3629 | 2149 |
| **% of Diff.methylated Enhancers** | **9.5** | **6.1** | **3.7** |
| Mutations in Enhancers | 1536 | 1536 | 1536 |
| Mutations in Diff.methylated Enhancers | 297 | 219 | 99 |
| **% of Mutations in Diff.methylated Enhancers** | **19.3** | **14.2** | **6.5** |
| **Fisher's Exact test (p-value)** | **2.64E-32** | **8.81E-31** | **9.0E-08** |

**Tables S6-S24- excel files**

**Table S1.** Steps in mutation filtering with the code upon mutation calling.

**Table S2.** Lack of correlation of mutations with open ATACseq peaks, see Fig. 2B as example.

**Table S3** Correspondence of mutations in enhancers to hypermethylation patterns in GDR and Osi treated cells.

**Table S4** Correspondence of mutations in enhancers to hypomethylation patterns in GDR and Osi treated cells.

**Table S5** Correspondence of mutations in active enhancers in HCT116 cells to the overall differentially methylated, hyper- and hypomethylation patterns in GDR cells.

**Tables S6-S21** VCF files after filtration for all drug and stress-resistant cells used in this study.

**Table S22** The list of differentially methylated sites in osimertinib-resistant cells

**Table S23** The list of differentially methylated sites in GDR cells

**Table S24** The list of ATACseq peaks in parental clone
